# Regulation of hippocampal excitatory synapse development by the adhesion G-protein coupled receptor Brain-specific angiogenesis inhibitor 2 (BAI2/ADGRB2)

**DOI:** 10.1101/2025.02.02.636169

**Authors:** Christina M. Meyer, Olga Vafaeva, Henry Low, David J. Speca, Elva Díaz

## Abstract

Glutamatergic synapses and their associated dendritic spines are critical information processing sites within the brain. Proper development of these specialized cellular junctions is important for normal brain functionality. Synaptic adhesion G protein-coupled receptors (aGPCRs) have been identified as regulators of synapse development and function. While two members of the Brain-specific angiogenesis inhibitor (BAI/ADGRB) subfamily of synaptic aGPCRs, BAI1/ADGRB1 and BAI3/ADGRB3, have been found to mediate synapse and spine formation, BAI2/ADGRB2 function remains uncharacterized at the synapse. Here, we show that endogenous BAI2 is expressed throughout the nervous system with prominent expression in synapse dense regions of the hippocampus. In dissociated hippocampal cultures, BAI2 is highly enriched at large postsynaptic sites, defined by the size of the postsynaptic scaffold PSD95. Loss of BAI2 negatively impacts glutamatergic synapses across development in dissociated hippocampal cultures. In contrast, GABAergic synapse density is unchanged. Furthermore, BAI2 deficient neurons have significant alterations in spine morphology with decreased density of mature PSD95-containing mushroom-shaped spines compared with wild-type neurons. Interestingly, no major alterations in dendritic complexity were observed in BAI2 deficient neurons, in contrast to previous results for the other BAIs. The reduction in mature mushroom-shaped spine is commensurate with a reduction in spine volume and head diameter. Altogether, these results demonstrate that the aGPCR BAI2 is an important regulator of glutamatergic synapse and PSD95-associated spine development in cultured hippocampal neurons. These results expand the knowledge of the BAI subfamily of aGPCRs in mediating excitatory synapse and spine development and highlight differences unique to BAI2.

**Highlights:**

- BAI2 expression is upregulated during postnatal development in the hippocampus
- BAI2 is enriched at postsynaptic sites, particularly large synapses
- Loss of BAI2 results in decreased density of glutamatergic synapses with no change in GABAergic synapse density
- Loss of BAI2 results in decreased density of mushroom spines

## INTRODUCTION

The proper development of synapses within the central nervous system relies on bi-directional signaling between the pre- and postsynaptic terminals. Trans-synaptic adhesion molecules (TSAMs) span the synaptic cleft and organize critical developmental events at the pre- and postsynaptic terminal (1), and a subset of TSAMs facilitate bidirectional signaling required during synaptogenesis (2–4).

The brain-specific angiogenesis inhibitor (BAI) subfamily of adhesion G-protein coupled receptors (aGPCRs) are a proposed TSAM group that facilitates synaptic development (3,5). The aGPCRs are characterized by an extensive extracellular N-terminus with multiple adhesion domains that facilitate cell-to-cell or cell-to-cell matrix interactions in addition to the classic seven transmembrane GPCR topology (6). Several aGPCR families, including the Latrophilins (ADGRL) (7) and the atypical CELSR/Flamingo cadherins (8), have known roles as TSAMs, suggesting a potential conserved role for neuronal aGPCRs. Mutations to the BAI proteins identified through whole exome sequencing are associated with neurodevelopmental and psychiatric conditions in human patients (9–11), though causative roles in any condition have not been elucidated.

While groups have identified BAI1 (12,13) and BAI3 (14–16) as critical promoters of glutamatergic synapse development, no such role in the developing nervous system has been reported for BAI2. Like the other BAI proteins, BAI2 contains a long extracellular N-terminal adhesive segment of over 900 amino acids, followed by a G-protein autoproteolysis (GAIN) domain, and the seven transmembrane structure with an intracellular C-terminus that defines GPCRs. BAI proteins also contain intracellular signaling motifs, including the extreme C-terminal QTEV PDZ-binding domain (PBD) (17). However, BAI2 appears to be functionally unique among the BAI proteins, as BAI2 does not bind to the N-terminal interaction partners identified for its relatives (18), and gain-of-function mutations are the putative cause of a rare spastic paraparesis condition in humans unique to BAI2 (19). Thus, we sought to investigate the role of BAI2 in synapse development.

Our initial interest in BAI2 arose from the results of a mouse forward mutagenesis screen in which we screened mutagenized animals for increased locomotor activity and mapped a locus for hyperactivity as a dominant quantitative trait locus (20). Whole exome sequencing uncovered a single point mutation in *Bai2* which results in a missense R619W mutation in the GAIN domain and reduced cell surface expression that likely impacts function. Notably, locomotor hyperactivity is unique to BAI2 among the BAI family, and a *Bai2* null mouse line (*Adgrb2^tm1b(KOMP)Mbp^*) tested by the International Mouse Phenotyping Consortium (IMPC) exhibits a similar hyperactivity phenotype seen in carriers of the *Bai2^R619W^* mutation that we identified.

Here, we characterize endogenous BAI2 protein expression during development in mouse brain and dissociated hippocampal neurons with a validated anti-BAI2 specific antibody. We show that BAI2 is highly enriched at large postsynaptic sites, defined by the size of the postsynaptic scaffold PSD95. We further demonstrate that BAI2 is important for vGLUT1-positive synapse and PSD95-positive spine development, with limited effects on dendritic arborization and morphology, in dissociated cultures of hippocampal neurons. In contrast, loss of BAI2 does not impact GABAergic synapse development. These results expand our understanding of the BAI subfamily of aGPCRs in mediating synapse and spine development and highlight differences unique to BAI2.

## MATERIALS AND METHODS

### Animals

All animal procedures followed National Institutes of Health (NIH) guidelines and have been approved by the Institutional Animal Care and Use Committee at the University of California, Davis. Animals were maintained at 12 light/12 dark cycle. Food and water were provided *ad libitum*.

The *Bai2* mutant allele (*Adgrb2^tm1b(KOMP)Mbp^*) was obtained from the Knockout Mouse Project (KOMP; www.komp.org) repository (21) and backcrossed 6 generations onto the C57BL/6J strain. Genotypes were determined with verified PCR primers (**Table 1**) with known samples as positive controls. Wild-type (WT) and knockout (KO) littermates of both sexes from heterozygous breeding pairs were used for experiments.

**Table 1.**
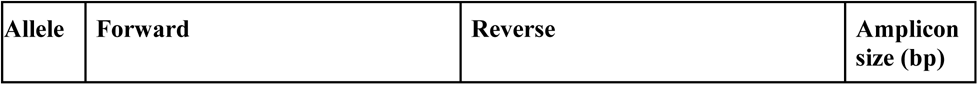
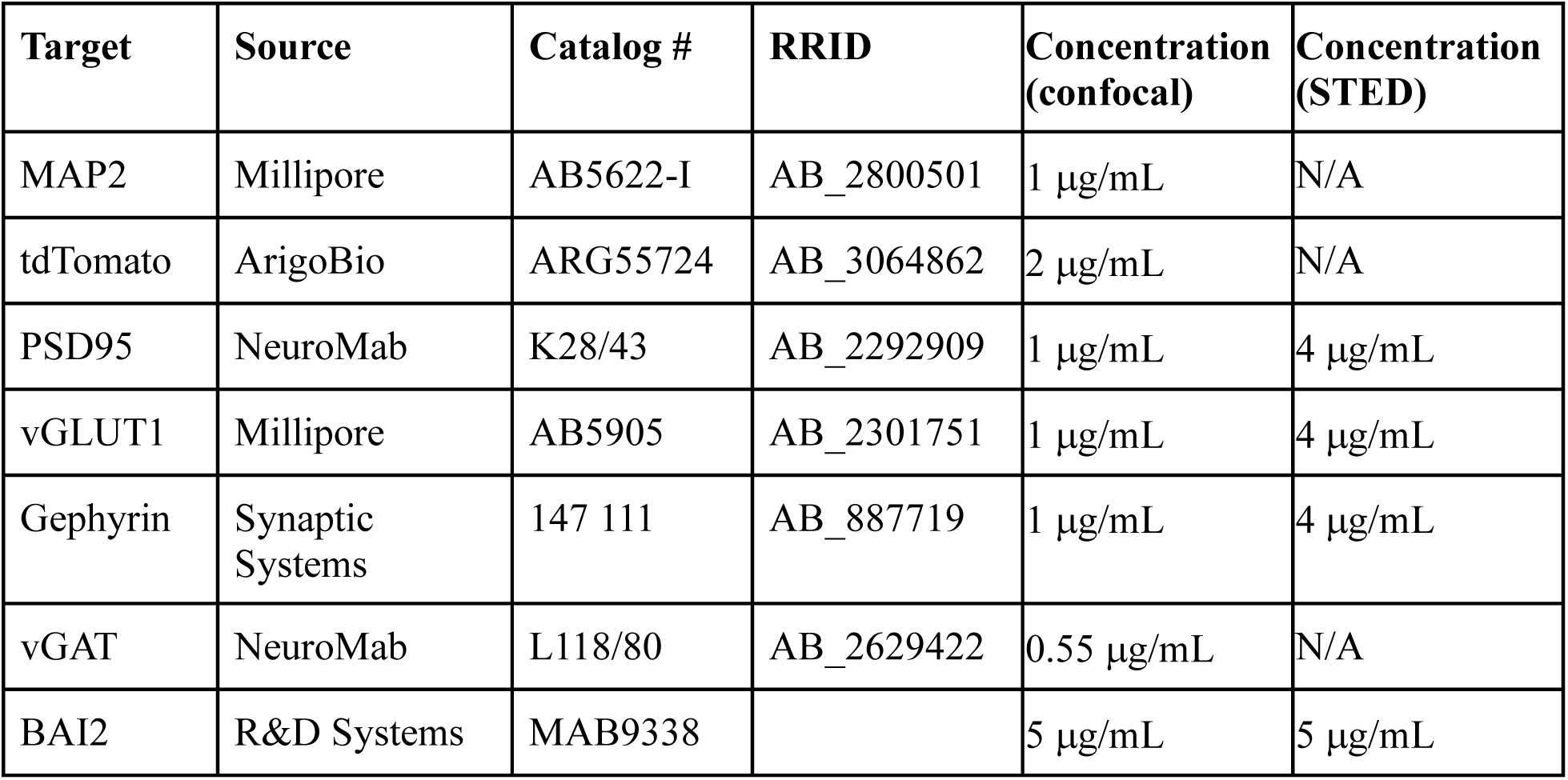
Primers for DNA genotyping.

### Dissociated hippocampal cultures

Primary mouse hippocampal neurons were plated over an astrocyte feeder layer (22). The astrocyte feeder layer was derived from postnatal day 1-2 (P1-2) rat cortex and grown to 70–90% confluency on collagen (Corning 354236) and poly-l-lysine (Sigma Aldrich P1399)-coated 25 cm^2^ plates in astrocyte plating medium consisting of 1X MEM (Gibco 11095080), 10% donor horse serum (Gibco 16050122), 0.6% D-glucose (Fisher D16-500; diluted from 30% stock made in 1X MEM), and 100 μg/mL penicillin/streptomycin (Gibco 15070063). The plates were shaken at 250 rotations per minute (RPM) overnight to remove other cell types from the culture. After shaking, purified astrocyte cultures were plated into 6-well plates (Fisher Scientific 07-200-83). Astrocytes were maintained in astrocyte plating medium until they reached 80-90% confluency. At 24 hours before neuronal plating, media was switched to Neuronal Maintenance Media (NMM) consisting of 1X Neurobasal (Gibco 21103049), 1X GlutaMAX (Gibco 35050061), and 1X B-27 supplement (Gibco 17504044).

Prior to hippocampal dissection, coverslips were treated with 1 M nitric acid for at least 12 hours, washed, and sterilized. Paraffin was heated to >90^°^C and 3-4 drops of melted paraffin were added to each coverslip to create a stage. Coverslips were then coated with 1 mg/mL poly-L-lysine (Sigma Aldrich P1399) diluted in borate buffer and incubated overnight at 37^°^C. The morning of dissection, coverslips were washed 3 times with distilled water and incubated for at least one hour with Neuronal Plating Media (NPM) consisting of 1X MEM (Gibco 11095080), 10% donor horse serum, 0.45% D-glucose, 1X sodium pyruvate (Gibco 11360070).

Primary mouse hippocampal neurons were cultured using previous methods (23,24), with minor modifications highlighted where relevant. Briefly, hippocampi were cultured from P0-1 WT and mutant mice obtained through heterozygous breeding and dissociated in Hank’s buffered saline solution (HBSS, Gibco 14025092) containing 2% filter-sterilized Papain (Roche 10108014001) for 12 minutes, quenched and triturated in NPM and plated at 0.6 - 0.65 × 10^5^ cells/well on coverslips in 6-well plates (Fisher Scientific 07-200-83). After 3-3.5 hours, coverslips were transferred and inverted over wells containing an astrocyte feeder layer and NMM. At 3 days *in vitro* (DIV), cultures were treated with 98% Cytarabine/Ara-C (Gibco 449560010) or floxuridine/FuDR (EMD Millipore F0503) at a final concentration of 2.5 μM to inhibit glial growth. A third volume change of the NMM was performed every 5-7 days, with weekly redosing of glial inhibitors. Hippocampal cultures were maintained for 13 or 17 DIV before being fixed and processed for immunocytochemistry. For lentiviral transduction, 1×10^9^ particles of adeno-associated virus (AAV9) expressing mCherry under the CMV promoter were delivered to the cell media at DIV7.

### Immunocytochemistry

For **confocal microscopy**, hippocampal cultures were fixed with 4% paraformaldehyde (PFA) with 3% sucrose for 12 minutes at room temperature (RT), rinsed 3 times in phosphate buffered saline (PBS) adjusted to pH 7.4, permeabilized in PBS with 0.02% Triton-X100 (PBST), and blocked with 5% IgG-free bovine serum albumin (BSA) for 60 minutes at RT. Antibodies were incubated overnight at the concentrations listed (**Table 3**) at 4°C in 1% BSA-PBST. Cells were washed with PBS for 3 times, 10 minutes each, blocked in 2.5% BSA-PBST for 30 minutes, and then incubated with secondary antibodies diluted in 1% BSA-PBST for 1 hour at RT. Following washes in PBS for 3 times, 10 minutes each, coverslips were mounted on microscope slides with ProLong Diamond mounting medium.

**Table 3.**
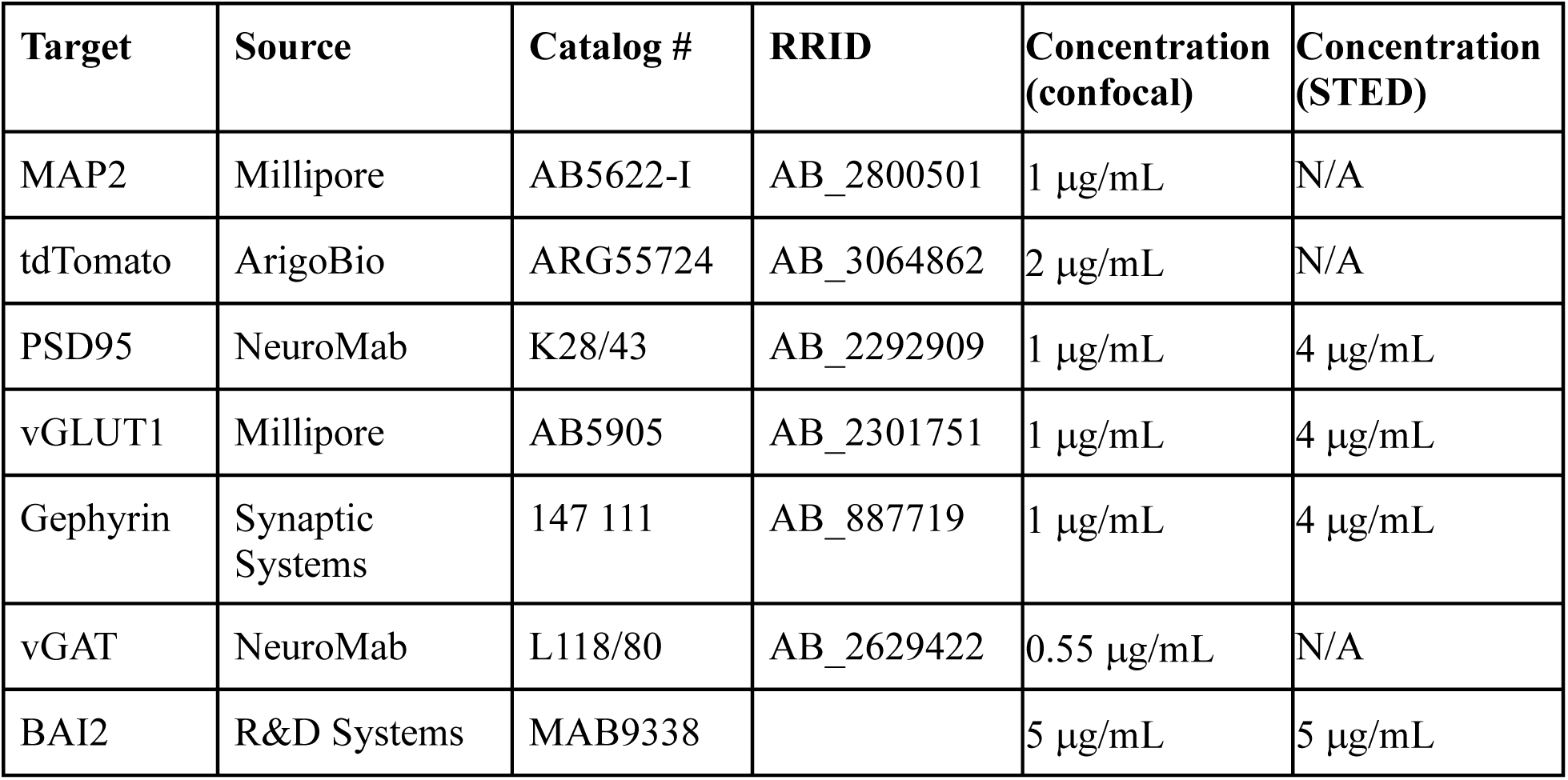
Antibodies and dilutions.

For **super-resolution microscopy**, hippocampal cultures were fixed with 3% glyoxal (EMD Millipore 128465) for 30 minutes at RT, quenched in 50 mM ammonium chloride for 20 minutes, rinsed 3 times in PBS, and processed as described above. For BAI2-labeled neurons, anti-BAI2 antibody is incubated with live neurons at RT for 10 minutes. After a brief rinse with PBS at RT, secondary antibody was incubated with live neurons for 15 minutes. Cells were then fixed and permeabilized as previously described.

### Immunohistochemistry

Mice (P7 to 3-months old) were deeply anesthetized with isoflurane and transcardially perfused with 4% PFA prepared fresh in 0.1 M sodium phosphate buffer (PB) with pH adjusted to 7.4. For P1 mice, brains were extracted and fixed in 4% PFA in 0.1 M PB overnight. After fixation brains were removed and cryoprotected with 30% sucrose solution. Cryopreserved tissue was sectioned with a sledge microtome equipped with a freezing stage and a series of 30 µm- thick tissue sections were collected in 0.1 M PB.

Immunohistochemistry was performed on free-floating or pre-mounted free-floating mouse brain sections prepared as described above. Immunohistochemistry with antibodies against BAI2 required antigen retrieval modification as follows: Prior to permeabilization, sections were treated with 0.05% Trypsin-EDTA for 20 minutes at 37°C. After incubation, the tissue slices were washed twice with 0.1M PB before proceeding with subsequent staining steps, listed below. Tissue sections were blocked with 10% donor horse serum in 0.1 M PB and 0.3% Triton for 1 hour at RT. Sections were then transferred to incubation buffer consisting of 5% normal horse serum in 0.1 M PB and 0.3% Triton with the primary antibody overnight at 4°C on an orbital shaker. After incubation, tissue was washed 3 times for 15 minutes in 0.1 M PB on an orbital shaker. Sections were then incubated with secondary antibody in the incubation buffer for 1 hour at RT on the shaker, protected from light. Brain sections were washed 3 times for 10 minutes in 0.1 M PB at RT on an orbital shaker and mounted on glass slides. After the mounted sections were dry, the slides were coated with ProLong Diamond mounting medium and cover slipped.

### Imaging and analysis

For **immunohistochemistry**, images were collected using a Leica TCS SP8 confocal microscope under a 20×/0.75 oil-immersion objective and stitched together in LAS X software. Fitted neuroanatomy maps were created using the BigWarp plugin for ImageJ/FIJI software (https://imagej.net/plugins/bigwarp), a tool for manual, landmark-based image alignment based on the anatomical outlines obtained from the online brain atlas (https://labs.gaidi.ca/mouse-brain-atlas/) developed by Dr. Matt Gaidica (Department of Neuroscience, Washington University, St. Louis, MI) based on atlas source (25).

For **immunocytochemistry**, images were acquired using a Leica SP8 instrument in either confocal (63× for synapses or 100× for spines) or Stimulated Emission Depletion (STED) mode (100×) under a 1.4 oil-immersion objective, and with constant settings for gain and offset between groups. Pinhole was set at 1 A.U. and resolution of 2048x2048 and a final resolution with 91.11 pixels/nm (confocal) and 22.18 pixels/nm (STED) were used for all images. Standard protocol during STED imaging involves imaging individual channels at consecutively higher intensity depletion to produce several “steps” of depletion, from no depletion to up to 4-5 levels of depletion (increasing the intensity of the depletion beam ∼5-10% each time). From there, all intensity plots across 10-20 individual puncta are mapped to one another to determine optimal imaging parameters.

For quantitative analysis of synapse density and puncta size, images were imported into FIJI/ImageJ software. All channels were thresholded to include all recognizable punctate structures in the analysis; to ensure consistency of and decrease for the thresholding process, 5-6 randomly chosen blinded images were selected for thresholding to determine appropriate thresholds for the dataset. 3 random stretches of dendrites (25-40 μm for vGLUT1, 30-60 μm for vGAT) representative for each cell were selected and cropped for synapse density analysis.

Multichannel, maximum projection of Z-stacked images were generated in ImageJ and a separate graphical user interface (GUI)-based tool (“ColocalVision”; publicly accessible on GitHub at: https://github.com/henrylowgh/ColocalVision) with customizable analysis parameters was developed using previously unpublished scripts to reduce analysis bias and efficiently standardize colocalization assessment procedures by facilitating threshold optimization, binary masking, and particle measurement. Scripts were written using the Groovy/IJM scripting languages and employed as an ImageJ plugin. Sufficient colocalization for a synapse was defined as any punctate structure with pixel-based channel overlap, which was checked manually and corresponded to an average of 5-20% pixel-based overlap of identified, thresholded pre- and post-synaptic markers. The resultant mask of colocalization was counted and processed to calculate synapse density per stretch, which was averaged per cell.

STED images were deconvoluted using Huygens Professional (Scientific Volume Imaging, ver 14). When using Huygens, a random sampling of 5 images across genotypes are used to set standard parameters for the group. Post-deconvolution, 2-3 random images are selected for comparison. For each image, we compare (i) raw images, (ii) Huygens-deconvoluted images, and (iii) application of a Gaussian blur with background subtraction in ImageJ/FIJI to the raw images. We ensure that the intensity is scalable between groups (ii) and (iii); compare the full width at half maximum (FWHM) readings between groups (i), (ii), and (iii); and there are no artificial aberrations produced from group (i) to (ii).

Imaris software (Oxford Instruments, ver 9.7) was used to quantify the intensity of BAI2 immunofluorescent signal, the average distance between BAI2 and vGLUT1/PSD95, the Sholl parameters, and the volume and length of spines. For the immunohistochemistry, a mask of the entire hippocampal area was created. Then, the intensity of BAI2 was calculated as a surface inside the hippocampal area mask. For the distance analysis, centroid positions of BAI2, vGLUT1, and PSD95 were created as spots. Positions of spots in 3-dimensional space were used to calculate the average distance between centroid positions. For the Sholl parameters, a maximum-projection intensity map was made from a surface created for MAP2. The dendrites filament tool was used to detect cell branching, dendrite width, and length. For spines, a random sampling of 5 images across genotypes were used to set standard parameters. A mask of the cell-fill (mCherry) was then created, and the filaments tool was used to create a rendering of both dendrites and individual spines.

### Quantification and statistical analysis

For all experiments, neurons in equivalently dense regions were selected for analysis. A minimum of 12-15 cells per condition were utilized based on findings from pilot experiments to ensure sufficient statistical power (ꞵ = 0.8); three coverslips per condition per experiment were used as technical replicates to ensure consistent plating density, health, and phenotypes across an experiment. Coverslips that display inadequate health (i.e., lower plating density, irregular processes, synaptic puncta far outside of norms established in previous cultures) were removed from analyses; data shown passed all quality control measures for health and robustness. Image acquisition and data analyses were performed blinded to condition and genotype. All experiments were performed with paired littermates, and experiments were repeated with a minimum of two to three independent experiments.

Graphs and statistical analyses were performed in Excel and GraphPad Prism (version 10.3.1). The UC Davis Health Clinical and Translational Science Center Biostatistics resource was utilized for help with statistics.

## RESULTS

### Validation of anti-BAI2 antibody and *Bai2* null transgenic line

We obtained the *Bai2* null reporter line (*Adgrb2^tm1b(KOMP)Mbp^*), referred to here as *Bai2* knockout (KO) mice, wherein critical exons 6-9 are replaced with a cassette containing the *Engrailed2* splice acceptor, an internal ribosome entry site and LacZ encoding β-galactosidase (β-gal) (**Fig. 1A**). The presence of the mutant allele was confirmed via PCR on genomic DNA (**Fig. 1B**, **Sup. Fig. 1A**). The activity of the β-gal reporter gene in the brain has been documented at 8 weeks by the IMPC (26), demonstrating that at least some mutant allele transcript – containing exons 1-5 followed by the cassette – could be expressed. This transcript has potential to generate a protein fragment of BAI2 containing the adhesive thrombospondin (TSP) domains TSP1 and TSP2 but not TSP3 and TSP4 that, if expressed, would not be tethered to the plasma membrane and lacks most of the functional domains of other aGPCRs. Thus, it is unlikely that this hypothetical fragment has biological activity; however, we cannot rule out this possibility conclusively. The Ensembl database (27) suggests that BAI1 and BAI3 have alternate transcripts that may encode proteins without the adhesive TSP domains, but no transcripts lacking the adhesive domains are reported for *Bai2*. Therefore, we do not anticipate that alternative variants of *Bai2* downstream of the LacZ cassette will be expressed from the mutant allele. However, further studies would be necessary to definitively assess possible alternative splice variants of *Bai2* that may be expressed during brain development.

**Figure 1.**
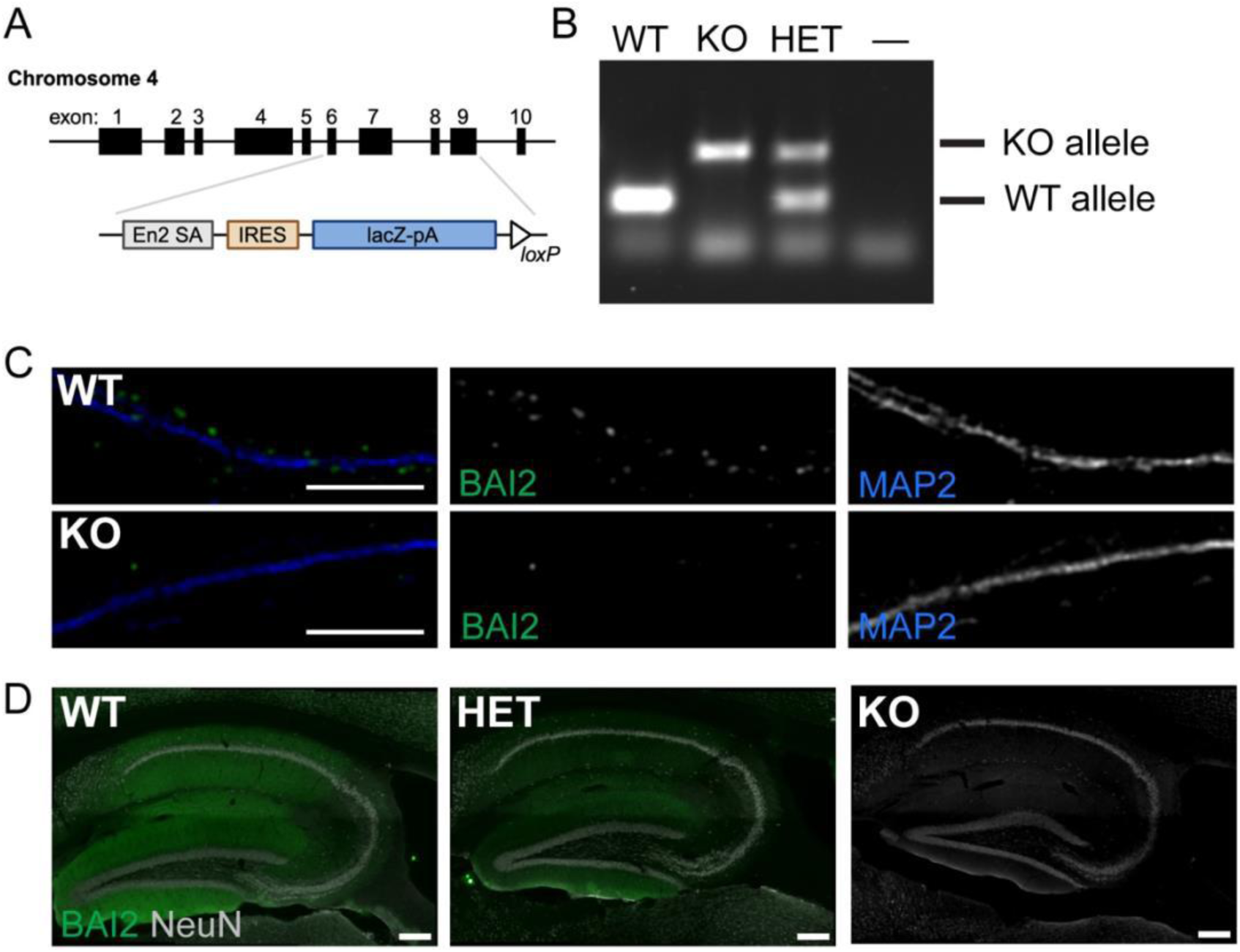
Validation of anti-BAI2 antibody with *Bai2* null mice. (A) Schematic of *Bai2* locus. In the *Bai2* null reporter line (*Adgrb2^tm1b(KOMP)Mbp^*), exons 6-9 are replaced with cassette containing an *Engrailed2* splice acceptor site (En2 SA) followed by an internal ribosome entry site (IRES) upstream of the *LacZ* gene. (B) Representative genotyping via polymerase chain reaction (PCR) to demonstrate presence of the wildtype (WT) and knock-out (KO) alleles. (C) Dissociated hippocampal neurons from WT and *Bai2* KO neurons stained with antibodies against BAI2 (green) and MAP2 (blue). (D) Brain sections from WT, heterozygous (HET), and *Bai2* KO 3-month-old mice stained with antibodies against BAI2 (green) and NeuN (white). Scale bar: 10 µm (C); 500 µm (D).

To define endogenous expression of BAI2 protein in brain, we optimized immunostaining using a commercially available anti-BAI2 antibody raised against the entire extracellular domain of BAI2. Using a series of deletion constructs, we determined that this anti-BAI2 antibody recognizes an epitope located within the TSP1-4 domains (**Sup. Fig. 1C, 1D**), which correspond to exons 4-7. Next, we validated the loss of BAI2 signal with immunocytochemistry (**Fig. 1C**) and immunohistochemistry (**Fig. 1D**) in dissociated hippocampal neurons and brain sections, respectively. Taken together, our results validate the loss of full-length BAI2 protein in the *Bai2* null mouse line and the specificity of the anti-BAI2 antibody.

### BAI2 expression in the mouse brain

We used the validated anti-BAI2 antibody to investigate BAI2 protein expression in the adult (P90) mouse brain using immunohistochemistry (**Fig. 2**). In the adult mouse brain, we determined that BAI2 has notable expression in the hippocampus, cortex, amygdala, striatum, and olfactory bulb (**Fig. 2A**). In the adult hippocampus, BAI2 is expressed in all CA regions, with the most pronounced staining in the molecular layers of the CA1 (**Fig. 2B**), a subregion critical for spatial and contextual memory (28). Within the CA1 field, BAI2 expression is found in the stratum lacunosum-moleculare, radiatum, and oriens, with a notable lack of detectable signal in the pyramidal layer, suggesting little to no expression on pyramidal cell bodies. BAI2 expression is also notable in the molecular layer of the dentate gyrus (DG, **Fig. 2B**), a region critical for episodic memory formation (29). Similarly, there was low level signal within the granule cell and hilus polymorphic layers, suggesting BAI2 is primarily expressed within neurites of granule cells. Expression in the CA3 region is much lower, present only within a subregion of the stratum oriens and radiatum (**Fig. 2B**). We conclude that BAI2 is primarily expressed in the neurites of pyramidal and granule cells in the hippocampus, though it is possible that other cell types also express BAI2 at a lower level.

**Figure 2.**
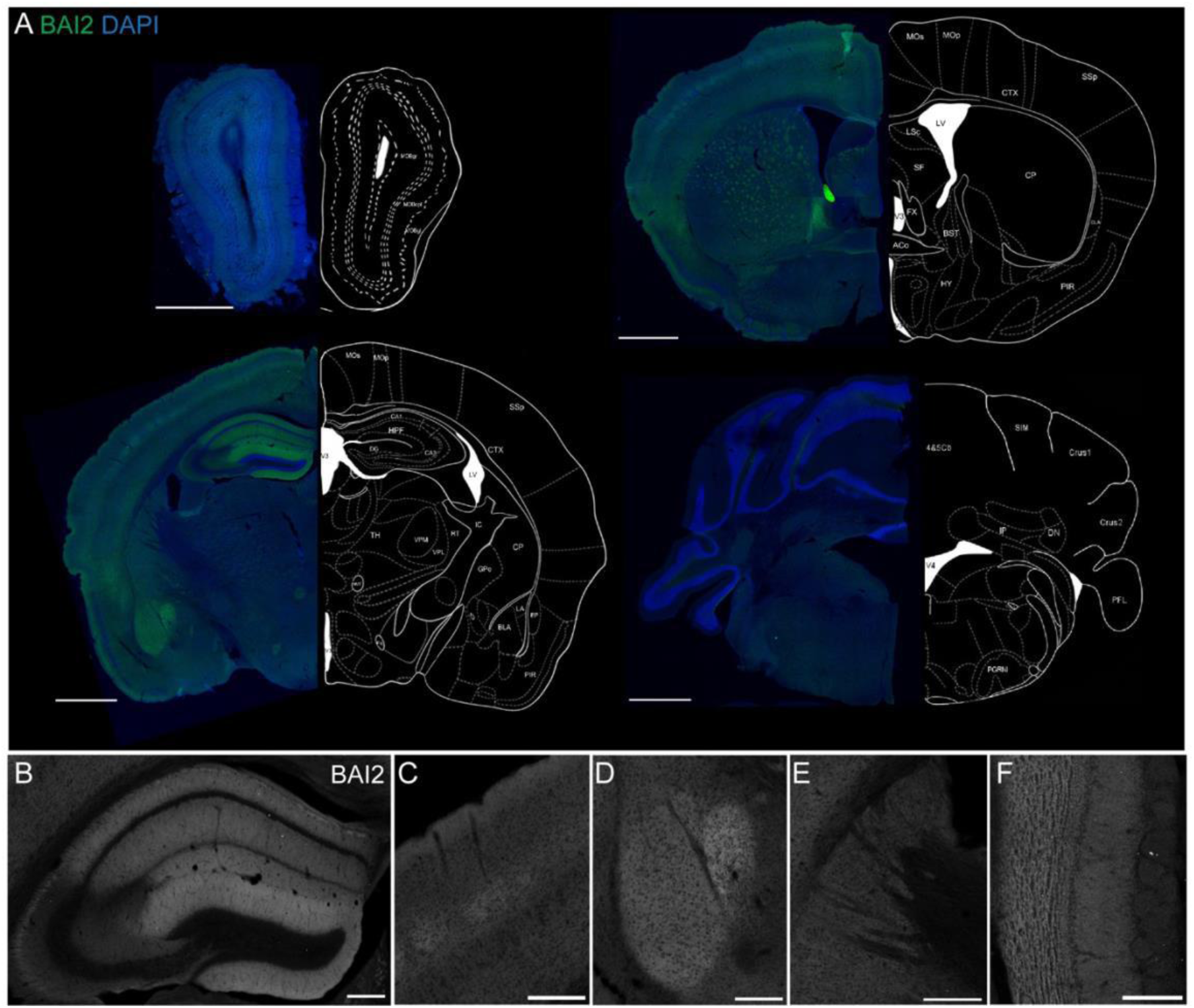
BAI2 expression in adult mouse brain. (A) Coronal sections of 3-month-old C57BL/6J wild-type mouse immunolabeled with validated anti-BAI2 antibody shows strong BAI2 protein expression in multiple brain regions. Insets show BAI2 expression patterns in hippocampus (B), cerebral cortex (C), amygdala (D), striatum (E), and olfactory bulb (F). Scale bar: 1 mm (A), 250 µm (B-F). Fitted neuroanatomy maps were created using the BigWarp plugin for ImageJ/FIJI software (https://imagej.net/plugins/bigwarp) based on the anatomical outlines obtained from the online brain atlas (https://labs.gaidi.ca/mouse-brain-atlas/). See Methods for details. Abbreviations: MOBgr: Main olfactory bulb, granule layer, MOBopl: Outer plexiform layer, MOBgl: Glomerular layer, ONL: Olfactory nerve layer, CP: Caudoputamen, ACB: Nucleus accumbens, PIR: Piriform area, EP: Endopiriform nucleus, ACO: Anterior commissure, MOp: Primary motor cortex, MOs: Secondary motor cortex, SSp: Primary somatosensory cortex, GPe: Globus pallidus, TH: Thalamus, BLA: Basolateral amygdala, LA: Lateral amygdala, IC: Internal capsule.

BAI2 was also expressed throughout the cerebral cortex, with highest apparent relative expression in layers I and IV (**Fig. 2C).** Layer I is composed of processes from several brain regions, including cortico-cortical and thalamic inputs and dendrites from cortical layers II through VIb (30), while layer IV is a main target and processing center for sensory signals from the thalamus (31). BAI2 expression is also notable in the amygdala (**Fig. 2D**), another structure in the temporal lobe associated with memory formation, as well as emotional and behavioral regulation such as fear conditioning and extinction (32,33). Within the amygdala, BAI2 staining was highest in the basolateral nuclei, the primary sensory input zone (34). We also found BAI2 within dendritic regions of the caudal dorsal striatum (**Fig. 2E**) (35). We did note patch-like expression of BAI2 within the rostral and intermediate caudoputamen (**Fig. 2A**), although the patches did not appear to correspond to the mosaic compartments known as the strisosomes (36). Further studies involving cell-specific markers would need to be performed to identify the specific cell populations expressing BAI2 in this region. And finally, in the olfactory bulb, BAI2 staining was strongest in the granule and outer plexiform layers, with weaker expression in the glomerular layer (**Fig. 2F**). Altogether, our results suggest BAI2 expression is likely highest in neurons, where it is primarily restricted to neurites.

### BAI2 expression is developmentally regulated in hippocampus

Based on the robust expression of BAI2 in the hippocampus, we focused our investigation of BAI2’s postnatal developmental time-course on this region. A previous study indicated that *Bai2* mRNA is developmentally regulated, with low expression in the embryonic brain and peak expression in the early postnatal period (37). To investigate BAI2 protein expression during development, we performed an immunohistochemical study ranging from P1 to P21 (**Fig. 3**). We observed that BAI2 expression was low in the first postnatal week, with more detectable expression at around P10 (**Fig. 3A-C**). At this stage, a low amount of BAI2 was detected in the stratum lacunosum-moleculare, the primary input site from the entorhinal cortex to the CA1 (38), as well as the molecular layer of the dentate gyrus, which contains the dendritic processes from granule cells. By P14, strong signal was detected in the CA1 and DG regions (**Fig. 3D**), which appears to be consistent with the expression pattern in the adult brain. BAI2 levels increased further throughout development, with the highest expression measured at P21 (**Fig. 3E, 3F**). Taken together, our observations confirm the expression of BAI2 in the hippocampal molecular layers during the period of hippocampal synaptogenesis.

**Figure 3.**
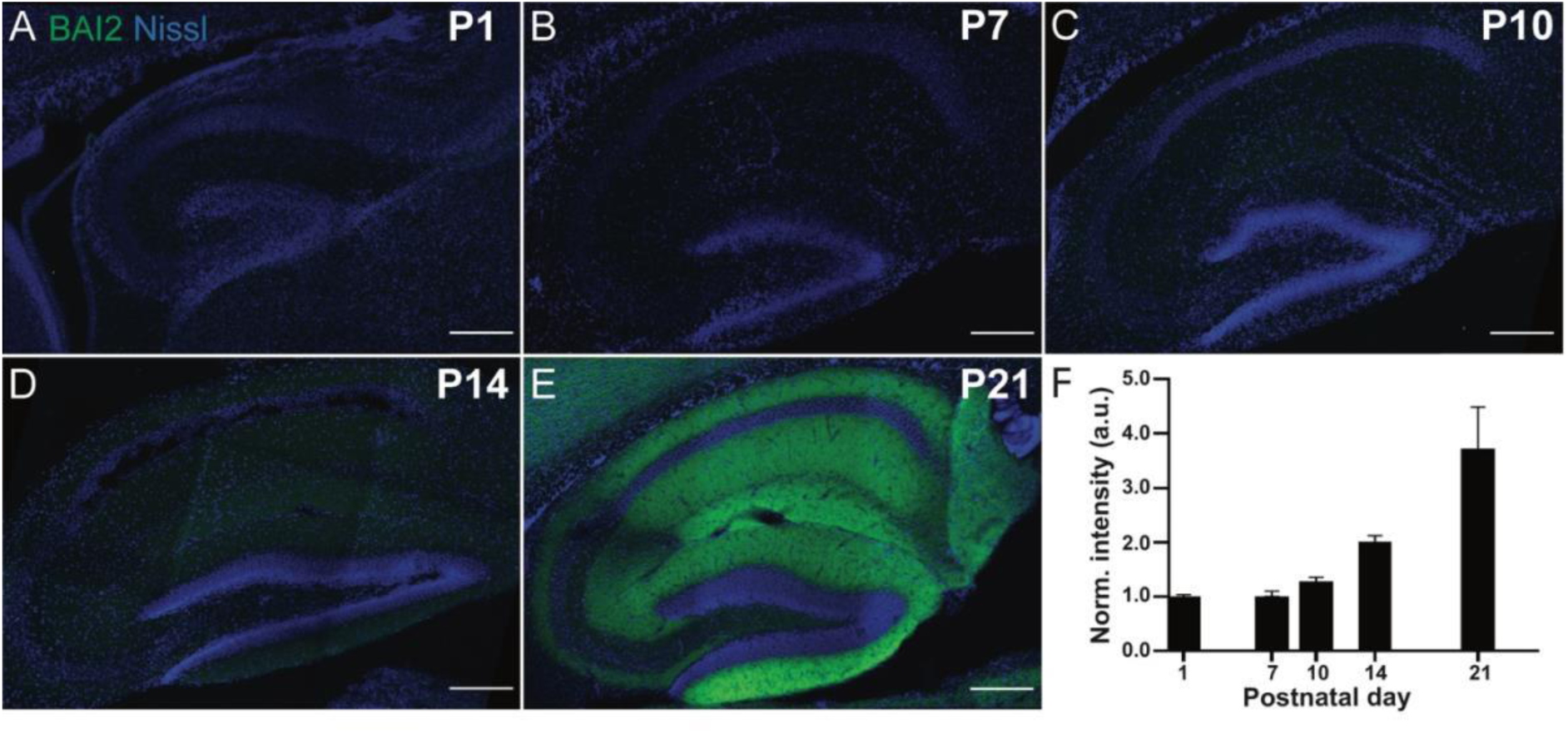
Developmental expression of BAI2 in mouse hippocampus. (A-E) Sagittal sections of mouse brain immunostained with anti-BAI2 antibody showing protein expression in the hippocampus at postnatal (P) day 1 (A), P7 (B), P10 (C), P14 (D), and P21 (E). Scale bar: 200 µm. (F) Graph depicts fluorescent intensity of BAI2 immunostaining in the hippocampus across the first three weeks of postnatal development. Data represent mean SEM of 5-8 sections per time point normalized to value at P1.

### BAI2 is localizes to larger postsynaptic sites at glutamatergic synapses

To determine BAI2 subcellular localization, we stained mature cultured hippocampal neurons with markers of glutamatergic synapses (**Fig. 4A**) and found that on average about half of glutamatergic synapses (defined as overlap of vGLUT1 and PSD95) contain BAI2 (**Fig. 4B**). We next sought to determine where BAI2 is localized at synapses. To further understand the relationship between BAI2 and pre- and postsynaptic structures, we utilized stimulated emission depletion (STED) super-resolution microscopy. We measured the average distance between BAI2 and markers of glutamatergic synapses, where we observed that the majority (>85%) of BAI2 signal was found within 1 micron of PSD95 and/or vGLUT1, with nearly all (>97.5%) being present within 3 microns of either marker **(Fig. 4C**). We also found that BAI2 appeared slightly closer to PSD95 than vGLUT1 **(Fig. 4C**), suggesting that BAI2 may be more closely associated with the postsynaptic terminal. However, it is important to note that the BAI2 N-terminus – where the anti-BAI2 antibody binds – is predicted to be in the synaptic cleft, making interpretation limited without a BAI2 C-terminal antibody for comparison. These data suggest that BAI2 has robust enrichment at and around glutamatergic synaptic sites.

**Figure 4.**
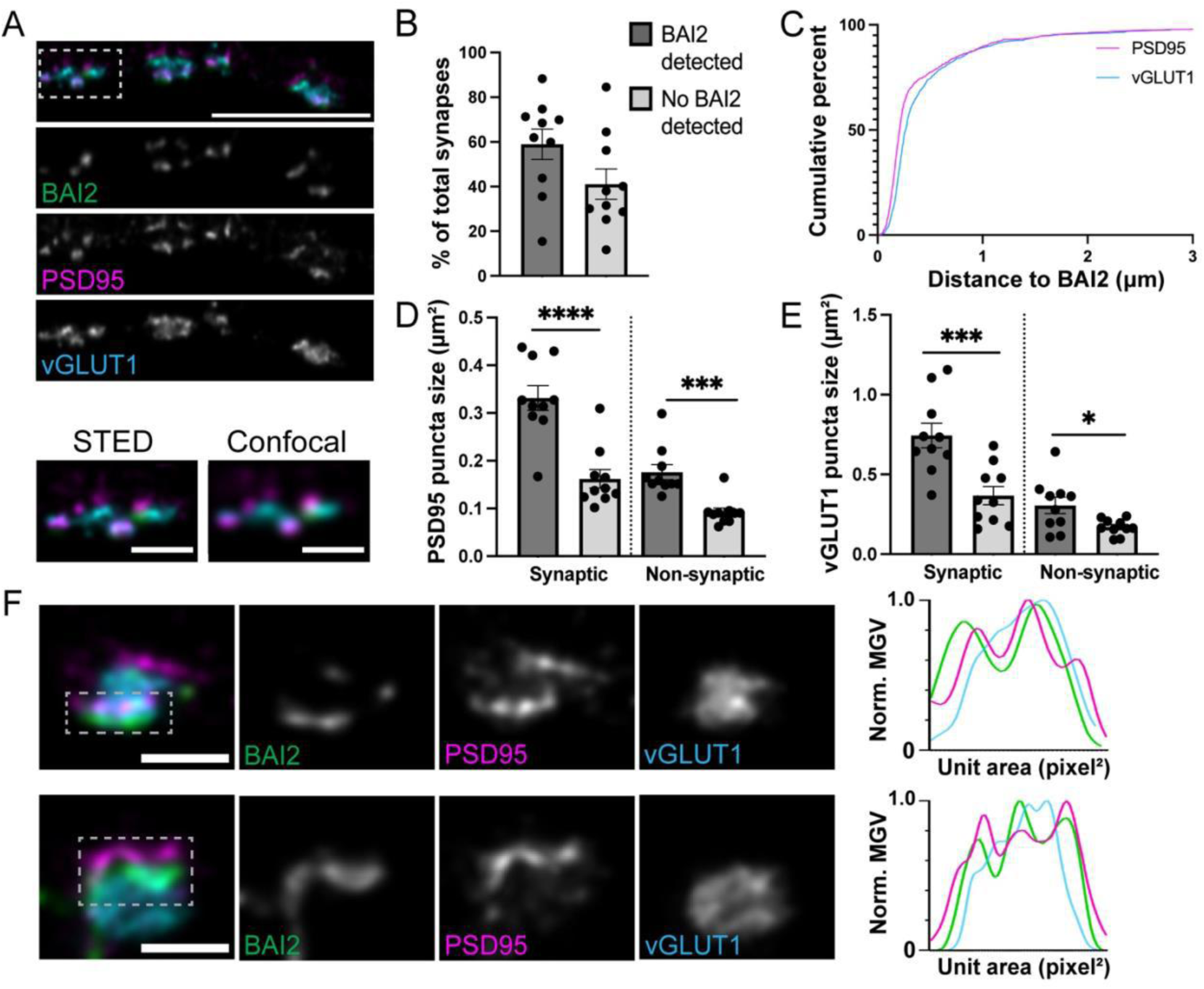
BAI2 is enriched at larger synapses. (A) Representative STED (Stimulated Emission Depletion) microscopy images of dissociated hippocampal neurons from wild-type mice at 17 DIV stained with antibodies against BAI2 (green), PSD95 (magenta), and vGLUT1 (cyan). Insets show STED mode and confocal mode for comparison. Scale bar: 10 µm (top), 2 µm (insets). (B) Graph depicts the percent of synapses (defined as overlap of vGLUT1 and PSD95 puncta) that contain BAI2 (dark gray bar) or do not contain BAI2 (light gray bar). (C) Graph depicts the cumulative histogram of the distance of BAI2 to vGLUT1 or PSD95. (D-E) Graphs depict the puncta size of PSD95 (D) or vGLUT1 (E) that overlap with BAI2 (dark gray bars) or do not overlap with BAI2 (light gray bars) at synaptic sites (defined as overlap of vGLUT1 and PSD95 puncta) or non-synaptic sites (defined as vGLUT1 or PSD95 puncta that do not overlap with PSD95 or vGLUT1, respectively). Data represent mean SEM. Statistics: t-test; *p<0.05, ***p<0.001, ****p<0.0001; n = 10 cells. (F) Representative higher magnification images demonstrating the nanoscale organization of BAI2 at individual synapses. The insets (dashed box) are shown as individual channels for BAI2 (green), PSD95 (magenta), and vGLUT1 (cyan). Graphs at right indicate the normalized mean gray value (MGV) of pixel intensity relative to pixel area. Note that BAI2 and PSD95 signals have a similar distribution in contrast to vGLUT1. Scale bar: 2 µm.

We stratified the size of vGLUT1 and PSD95 puncta from BAI2-positive and BAI2-negative synapses in WT neurons and found that the presence of BAI2 at a synapse is correlated with larger vGLUT1 and PSD95 puncta area (**Fig. 4D, 4E**). A similar analysis performed with non-synaptic puncta – vGLUT1 and PSD95 puncta that do not colocalize with the corresponding pre- or postsynaptic signal – found that the presence of BAI2 was also correlated with larger puncta area (**Fig. 4D, 4E**). Interestingly, we also found that BAI2 is organized into nanoscale structures at excitatory synapses, closely following the nanoscale organization of PSD95 as measured by mean gray volume over selected area (**Fig. 4F**). Further studies with proteins known to form nanocolumns that align across the pre- and postsynaptic terminals, such as RIM1/2 and GluA1, would be needed to further identify if BAI2 aligns with nanodomains at excitatory synapses. These results provide evidence that BAI2, like its relatives, is enriched in large postsynaptic terminals, and organizes with postsynaptic scaffolds such as PSD95.

### Loss of BAI2 results in decreased glutamatergic synapses without a strong effect on neuronal morphology

Previous studies demonstrated that loss of BAI1 results in dramatic alterations to neuronal morphology (39). To investigate effects on neuronal morphology due to loss of BAI2, we prepared hippocampal cultures from *Bai2* WT and KO littermates at two time points (DIV13 and DIV17) and immunostained with antibodies against MAP2 (**Fig. 5A**). In contrast to other BAIs, we observed minor changes to dendritic branching complexity (**Fig. 5B, 5C**) as assessed via Sholl analysis (40). There was also no change in dendrite length or thickness between *Bai2* WT and KO neurons (**Fig. 5D, 5E**). Thus, loss of BAI2 does not appear to play a critical role in neuronal morphology *in vitro*, including dendritic arborization and development.

**Figure 5.**
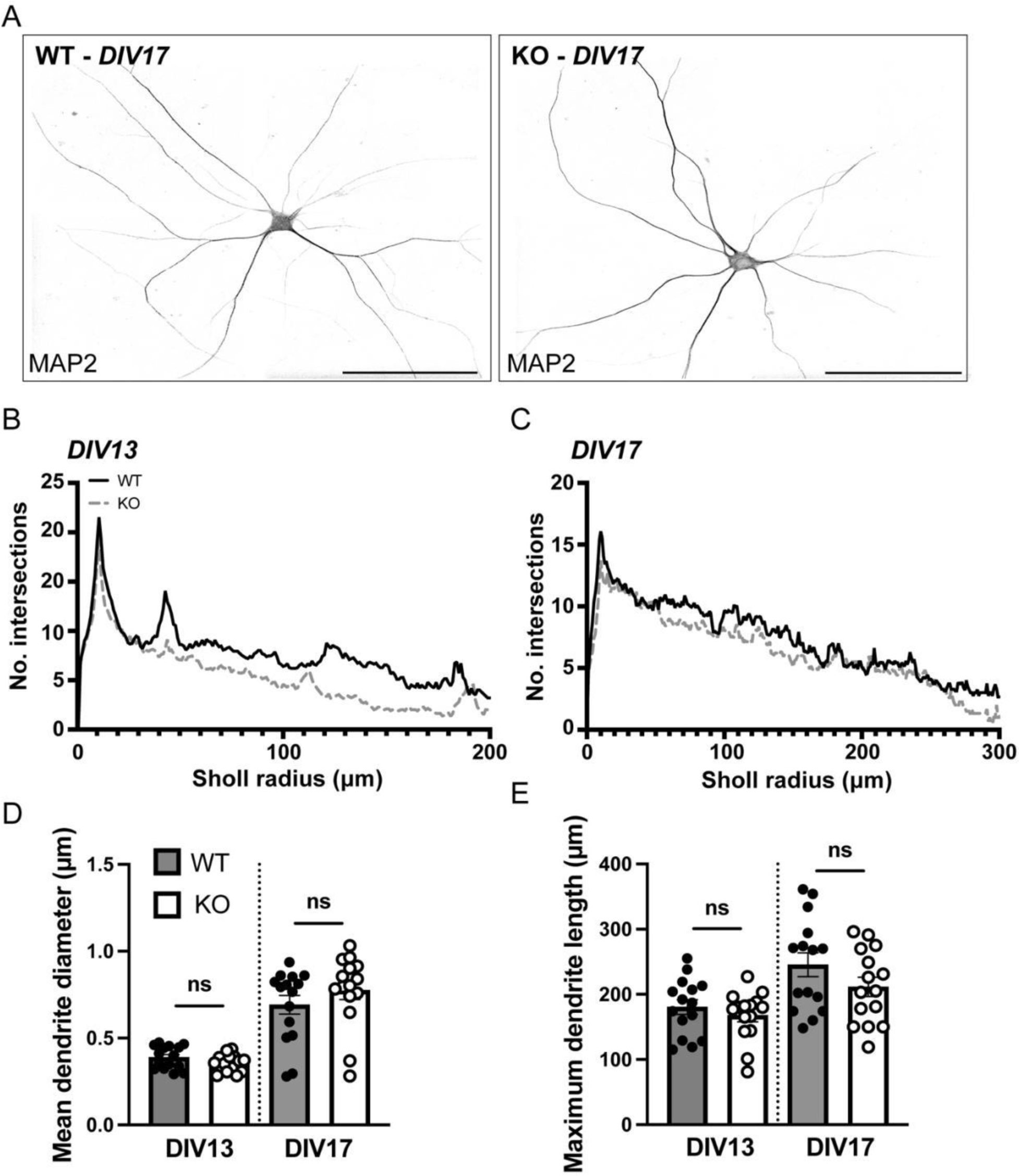
Dendritic branch length and thickness is unchanged in BAI2 deficient neurons. (A) Representative images of hippocampal neurons from wild-type (WT) and *Bai2* knockout (KO) mice at DIV17 immunostained with MAP2. (B-D) Graphs depict Sholl analysis of dendritic complexity (B), dendritic length (C), dendritic diameter (D), and maximum dendrite length (E) from WT and KO neurons at DIV13 and DIV17. Data represent mean±SEM. Statistics: t-test, ns, not significant; n = 15 cells/group. Scale bar 200 µm.

To characterize the effect of loss of BAI2 on synapse development, we generated hippocampal cultures from *Bai2* WT and KO littermates at two time points (DIV13 and DIV17) and used an image analysis pipeline to identify glutamatergic (**Fig. 6A, Sup. Fig. 2A**) and GABAergic (**Fig. 6C, Sup. Fig. 2C**) synapses with confocal microscopy. We found reduced number of glutamatergic synapses (defined as overlap of vGLUT1 and PSD95) along secondary and tertiary processes at both early (DIV13) (**Fig. 6B**) and late (DIV17) time points (**Sup. Fig. 2B**). In contrast, we did not observe any changes to GABAergic inhibitory synapse (defined by the overlap of vGAT and Gephyrin) density at either time point (**Fig. 6E, Sup. Fig. 2C**). Altogether, we conclude that BAI2 plays a role in the development of glutamatergic – but not GABAergic – synapses *in vitro*.

**Figure 6.**
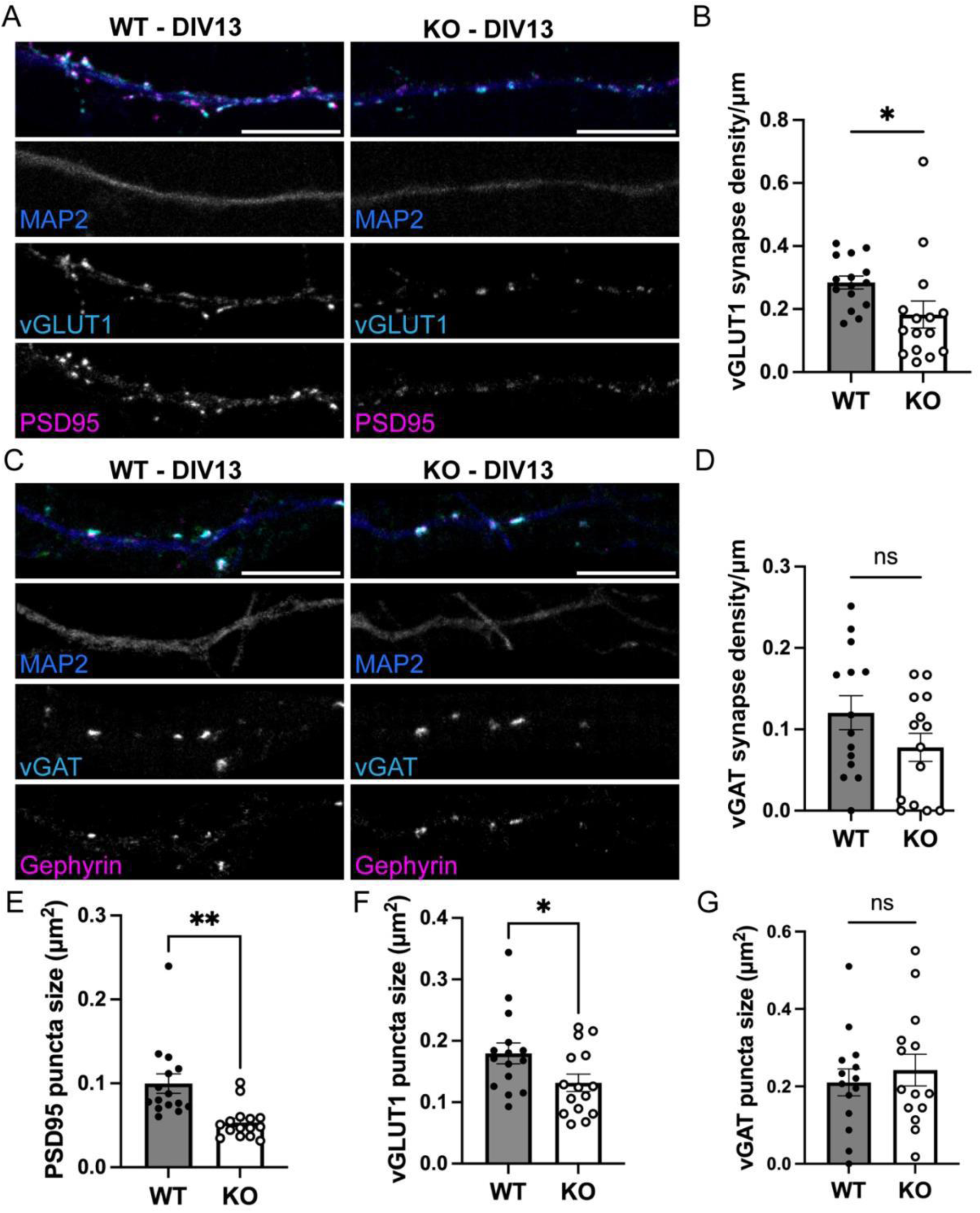
Loss of BAI2 results in reduced excitatory synapse density. (A, C) Representative images of hippocampal neurons from wild-type (WT) and *Bai2* knockout (KO) mice at DIV13 immunostained with presynaptic markers (vGLUT1 or vGAT) and postsynaptic markers (PSD95 or gephyrin). (B, D) Graphs depict density of vGLUT1-positive (E) and vGAT-positive synapses (D) at DIV13. (E, F, G) Graphs depict puncta size for PSD95 (E), vGLUT1 (F), and vGAT (G) at DIV13. Data represent mean±SEM. Statistics: unpaired t-test with or without Welch’s correction, ns, not significant, p*<0.05, p**<0.01; n = 14-16 cells/group. Scale bar: 10 µm.

We noted that BAI2 KO neurons had smaller average vGLUT1 and PSD95 puncta sizes at DIV13 (**Fig. 6E, 6F**), with no overall change in vGAT puncta size (**Fig. 6G**). However, we did not observe any consistent trend when comparing WT and KO puncta sizes at DIV17 (**S. Fig. 2E-G**), suggesting that the synapse density deficit cannot be solely explained by smaller synaptic puncta leading to less overlapping signals. Altogether, we conclude that BAI2 plays a role in the development of glutamatergic but not GABAergic synapses.

### Loss of BAI2 causes aberrations to spine development

Dendritic spines, actin-rich membrane protrusions along dendritic shafts, are the major receiving sites of excitatory synaptic input (41). To investigate alterations in spine development in the absence of BAI2, hippocampal neurons were transduced with a lentiviral vector expressing mCherry as a cell fill (**Fig. 7A**). Given BAI2’s relative enrichment in PSD95-positive synapses, we restricted our spine analysis to spines that contained PSD95. Interestingly, the distribution of spine morphologies (mushroom, stubby, filopodia) is different in *Bai2* KO neurons with reduced numbers of mushroom spines and increased numbers of stubby and filopodia spines (**Fig. 7B**). We observed that *Bai2* KO neurons display reduced density of mature mushroom spines and a concomitant increase in the density of more immature filopodial spines (**Fig. 7C**). This finding was also reflected in morphological features of spines in *Bai2* KO neurons, wherein we found an overall decrease in total spine volume, spine head size, and spine length (**Fig. 7D-7F**). These data demonstrate that BAI2 is necessary for proper PSD95-associated spine development in hippocampal cultures.

**Figure 7.**
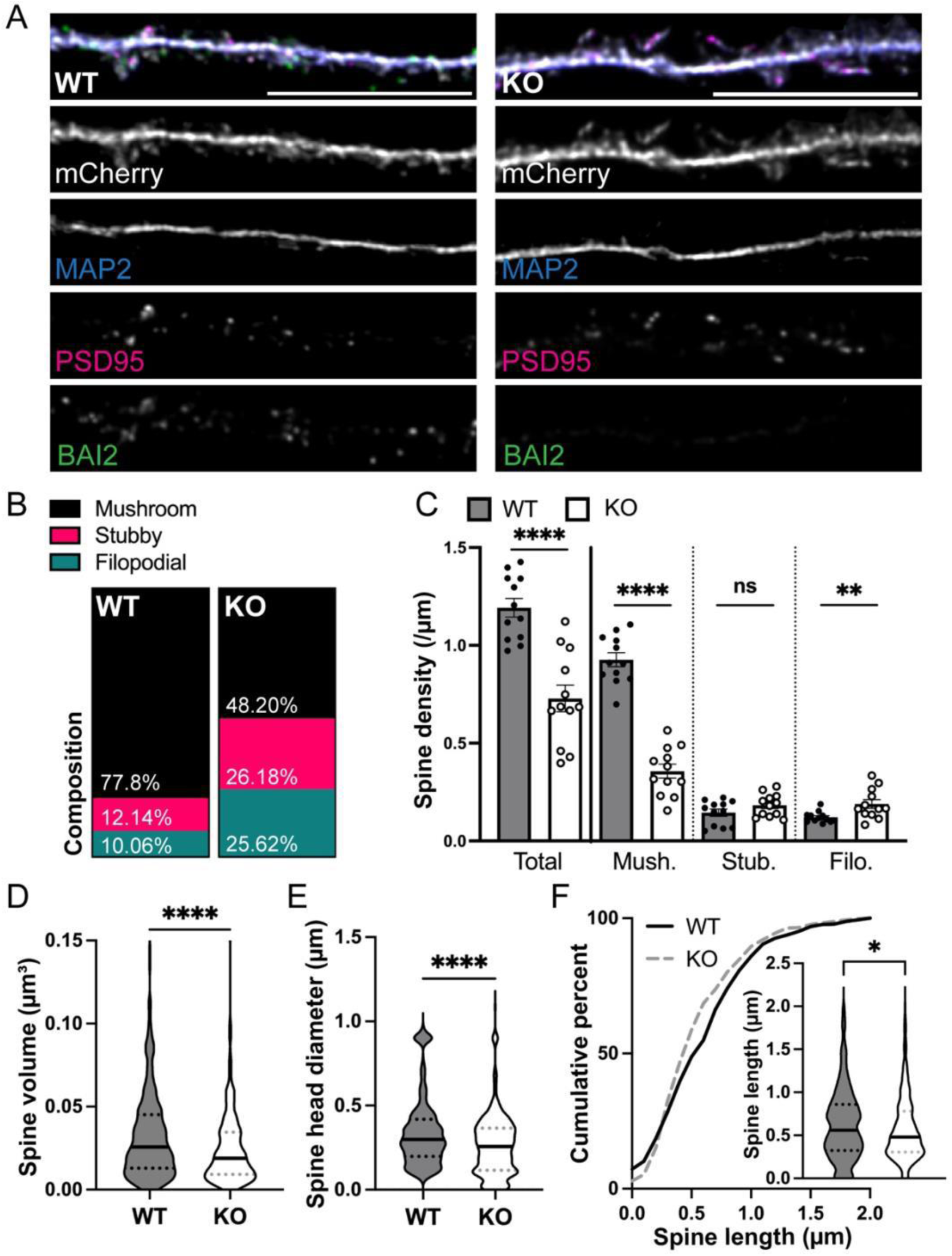
Loss of BAI2 alters distribution of spine morphological types. (A) Representative images of mCherry-transfected hippocampal neurons from wild-type (WT) and *Bai2* knockout (KO) mice at DIV17-8 and immunostained with antibodies against MAP2, mCherry (utilizing an anti-tdTomato antibody), PSD95, and BAI2. (B) Stacked bar graphs depict the average distribution of mushroom, stubby and filopodial type spines in *Bai2* KO and WT neurons. (C-F) Graphs depict density of mushroom, stubby, and filopodia spines (C), spine volume (D), spine head diameter (E) and spine length (F) in *Bai2* KO and WT neurons. Data represent mean±SEM Statistics: unpaired t-test, p*< 0.05, p**<0.01, p****<0.0001, ns, not significant; n = 12 cells, >500 spines/group. Scale bar: 10 µm.

## DISCUSSION

Members of the BAI aGPCR subfamily play important roles in neuronal development with distinct and overlapping functions. While numerous groups have identified BAI1 (12,18,42,43) and BAI3 (14–16) as critical for synapse development, the function of BAI2 (37,44) at synapses has remained relatively enigmatic (5). Furthermore, BAI2 appears at least somewhat functionally unique within the BAI family, as it does not bind the presynaptic binding partner identified for BAI1 and BAI3 (18), and mutations to BAI2 and not to any other BAI are causative of at least two motor phenotypes – hyperactivity (20) and progressive spastic paraparesis (19).

To address this gap, we explored the expression of BAI2 in the WT mouse brain, as well as the role of BAI2 in synapse and spine development using a *Bai2* deficient line obtained from the NIH-sponsored KOMP. While it is a formal possibility that a small fragment of the BAI2 protein is generated from the mutant allele transcript, it would not be associated with the plasma membrane of synapses and thus unlikely to exert any function. Thus, we interpret our phenotypic data as the effect of a loss of full-length BAI2 within mouse hippocampal neurons.

The optimization and validation of an anti-BAI2 antibody that recognizes endogenous BAI2 was critical for this study. With this tool, we observed that BAI2 is expressed in multiple regions in the adult mouse cerebrum, including the hippocampus, cortical layers, amygdala, striatum, and olfactory bulb. We noted a lack of signal in the cerebellum, thalamus, and hypothalamus, though it is possible that BAI2 is expressed at a much lower level in these regions. We were intrigued to see prominent expression of BAI2 within the CA1 and DG subfields of the hippocampus, as well as in other regions associated with decision-making, emotional processing, memory, and learning, such as the cortex, striatum, and amygdala. Because we found the most prominent expression and developmental regulation in the hippocampus and could reproducibly document BAI2 expression in hippocampal pyramidal neurons, we focused our subsequent studies on the function of BAI2 in dissociated cultures of hippocampal pyramidal neurons.

Within dissociated hippocampal cultures, we observed that BAI2 is present at approximately half of all excitatory synapses (defined by the overlap of vGLUT1 and PSD95). Approximately 60% of BAI2 was colocalized with glutamatergic synaptic markers in WT cells under super-resolution microscopy, with a majority (>85%) of BAI2 being localized within a micron of vGLUT1 and/or PSD95. Given that super-resolution microscopy leads to significantly less overlap of synaptic signals, we interpreted these findings as evidence that BAI2 is strongly associated with excitatory synapses. We also noted that BAI2 was commonly associated with larger postsynaptic sites, though it is possible that BAI2 may still be present at smaller synapses at a sub-detectable threshold. It would be interesting to determine if BAI1 and BAI3 are found in the same synapses as BAI2, or if each BAI protein labels a specific subset of synapses. If different combinations of BAI proteins were located at synapses, they could be a factor for synaptic specificity during the synaptogenesis phase, recruiting specific types of connections through binding different presynaptic binding partners. Alternatively, they could define other properties of excitatory synapses, such as their size, plasticity, or electrophysiological responses as metabotropic and mechanosensitive receptors, which has been proposed for aGPCRs more broadly (45).

We also compared cell morphology of dissociated neuronal cultures at DIV13 and DIV17, as loss of BAI1 (39) and BAI3 (10) have been shown to negatively affect dendritic branching, thickness, and development. To our surprise, we did not find dramatic effects on neuronal morphology in *Bai2* KO neurons. Though we did observe a minor effect on dendritic branching at DIV13, this difference was not apparent at later developmental stages, indicating that BAI2 plays a relatively minor role in dendritic branching at a specific developmental point. We also did not find a significant effect on dendrite morphology, as measured by mean dendrite diameter and maximum dendritic length. Thus, we conclude that BAI2 likely has a limited role in dendritic development in cultured neurons compared to BAI1 and BAI3.

Next, we demonstrate that *Bai2* KO neurons have a glutamatergic synapse density deficit, as measured by the overlap of vGLUT1 and PSD95, within dissociated cultures. This phenotype was observed at both DIV13 and DIV17, suggesting that the phenotype persists across the synaptogenesis period. At DIV13, dendrites with equal area and morphology were chosen for analysis to account for any apparent potential effect from the slight difference observed in dendritic branching. This control, in parallel with the apparent lack of significant changes in dendritic branching at DIV17, suggests that the glutamatergic synaptic phenotypes we have observed are independent of dendritic changes. Furthermore, we observed a decrease in the size of vGLUT1 and PSD95 at DIV13, but no consistent change in vGLUT1 and PSD95 puncta size at DIV17. In contrast, we did not see an effect on GABAergic synapse density or vGAT puncta size at either time point, suggesting that BAI2’s synaptic phenotype is specific for excitatory synapses within the dissociated hippocampal system. We have considered two possibilities: (1) that fewer vGLUT1 synapses are formed globally, and/or (2) that vGLUT1 synapses are less stable, and thus at the fixed timepoints we selected, we observe fewer synaptic terminals. Future studies involving live imaging could be utilized to assess changes in excitatory synapse density and the stability of excitatory synapses over time.

We also assessed spine morphology in DIV17-18 dissociated cultures transduced with mCherry. Given BAI2’s effects on PSD95, we restricted our analysis to spines that contained PSD95 somewhere within their volume. Intriguingly, we found a large change in PSD95-positive spine density, with a striking loss of mushroom spines containing PSD95 in the *Bai2* KO condition compared to WT, and concomitant increases in the relative abundance of stubby and filopodial-type spines containing PSD95. We were intrigued by the fact that the spine phenotype was more dramatic than the synaptic phenotype we observed. Given the differences between synapse and spine density, we hypothesize that more synapses may be forming on the dendritic shaft (46,47) in *Bai2* KO neurons compared to WT. Future studies are needed to determine the functional impact of the change in distribution from spine synapses to shaft synapses in *Bai2* KO neurons.

Taken together, our data suggests that BAI2 facilitates excitatory synapse and spine maturation. This phenotype would not be wholly unexpected of a BAI protein, as BAI1 has been identified as a critical protein for the clustering of AMPA-type receptors (AMPARs) at the PSD of cochlear ribbon synapses (48). Delivery and clustering of AMPARs at nascent synapses represents a critical step in later synapse development, indicating that BAI proteins may promote the formation of mature synapses. Alternatively, BAI2 may be recruited to maturing synapses, while maturation is driven by other TSAMs. Investigating the time course of BAI2 delivery to synapses in living neurons may help to distinguish whether BAI2 plays a direct role in synapse and spine maturation or is simply correlated with maturation.

More broadly, the extent to which BAI proteins are functionally unique remains unclear. Though all members appear to influence excitatory synapse density, there are significant divergences between BAI2 and other BAI proteins, including the lack of a binding interaction between RTN4R (18) and motor phenotypes (19,20). Thus, we expect that BAI proteins may not be fully redundant and mediate different aspects of excitatory synapse development. If the BAI proteins are not fully compensatory, we would expect to observe additive effects on synapse density in hippocampal neurons lacking two or more BAI proteins. Furthermore, we would expect that reintroduction of BAI2, but not BAI1 or BAI3, would rescue the synapse and spine deficits we observed if BAI2 acts independently of other BAI proteins. Future studies with structure-function and *Bai2* KO rescue approaches with modified forms of BAI2, as well as WT BAI1 and BAI3, will begin to address these outstanding questions.

In summary, we demonstrate a role for the aGPCR BAI2 in the regulation of excitatory synapse and spine development in hippocampal pyramidal cells for the first time. These results contribute to our understanding of the BAI subfamily of aGPCRs in mediating synapse and spine development.

## DATA AVAILABILITY STATEMENT

The raw data supporting the conclusions of this article will be made available by the authors, without undue reservation.

## ETHICS STATEMENT

The animal study was approved by Institutional Animal Care and Use Committee of the University of California, Davis. The study was conducted in accordance with the local legislation and institutional requirements.

## AUTHOR CONTRIBUTIONS

CMM, OV, DJS and ED: methodology and writing. CMM, OV: experimentation. CMM, OV, DJS and ED: formal analysis. CMM and HL: script writing and optimization. DJS and ED: funding acquisition and supervision. All authors reviewed the article and approved the final submitted version.

## ACKNOWLEDGEMENTS

We thank Drs. Kim McAllister, Jennifer Whistler, Karen Zito and members of the Diaz, Gray, Hell and Zito labs (University of California, Davis) for their insightful and valuable input and comments throughout the project. We also thank Drs. Hwai-Jong Cheng (Academia Sinica) and Karl Murray (University of California, Davis) for their expertise and input on the neuroanatomical studies and techniques. We are especially grateful to Dr. Ingrid Brust-Mascher (Advanced Imaging Facility, University of California, Davis School of Veterinary Medicine) for her expert assistance with microscopy and analysis. Additionally, we thank Drs. James Trimmer and Jody Martin (University of California, Davis) for generously sharing antibodies and lentiviral vectors, respectively, used in this research.

