## Supplemental Figures for "Regulation of hippocampal excitatory synapse development by the adhesion G-protein coupled receptor Brain-specific angiogenesis inhibitor 2 (BAI2/ADGRB2)"

**SUPPLEMENTAL FIGURES AND LEGENDS**

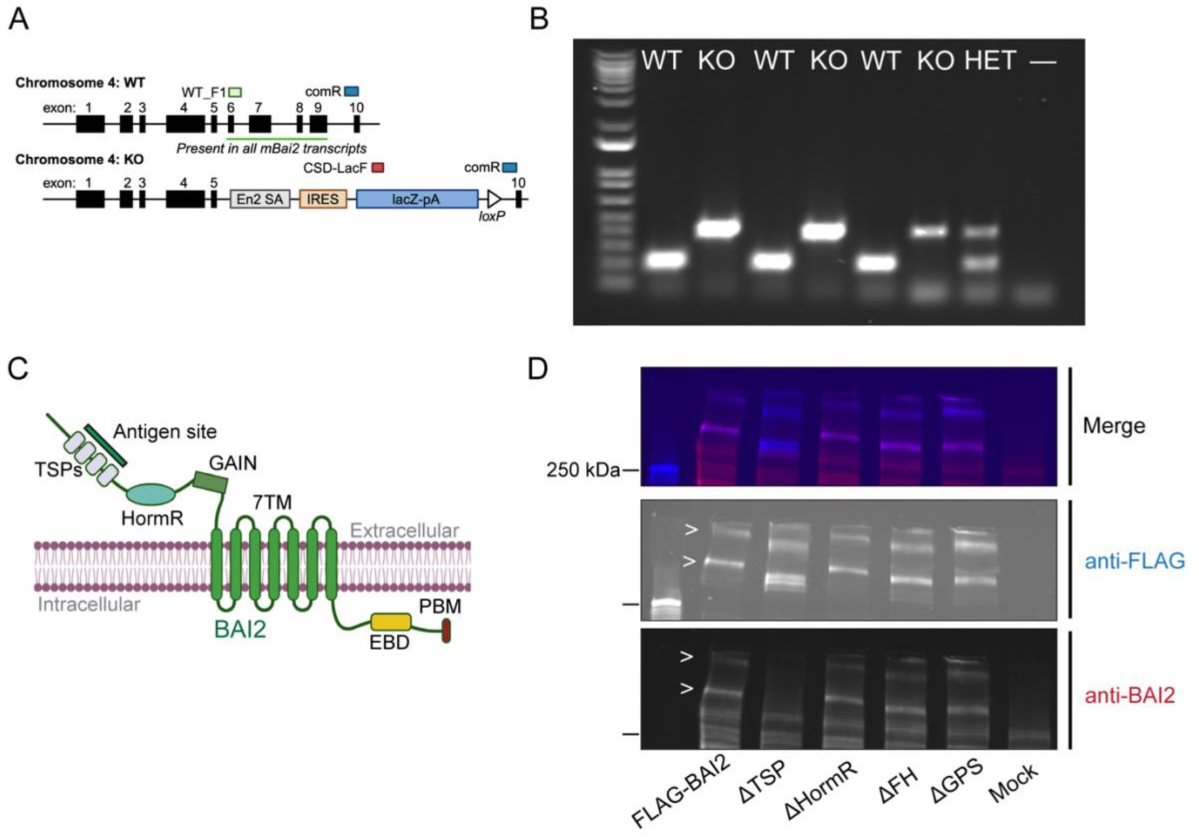


**Supplemental Figure 1. Additional information on *Bai2* null mice and antibody validation** (A) Schematic of *Bai2* wild-type (WT) and knockout (KO) alleles. Approximate locations of primer binding sites (WT_F1, CSD-LacF, and comR) are shown. (B) Full gel from Fig. 1B with increased contrast and multiple WT and KO samples from paired littermates. (C) Schematic of BAI2, labeled with relevant motifs. TSP: Thrombospondin type-1 repeat; HormR: Hormone binding domain; GAIN: GPCR autoproteolysis-inducing domain; 7TM: 7 transmembrane domain; EBD: ELMO binding domain; PBM: PDZ binding motif. The anti-BAI2 antibody binding epitope at the TSP repeats is labeled. (D) Representative immunoblot for antibody epitope mapping. FLAG-tagged BAI2 (FLAG-BAI2) constructs, consisting of the full-length protein and a series of deletion mutants of the previously listed domains, were expressed in HEK293T cells. Lysates were probed with anti-FLAG and anti-BAI2 antibodies. The high-molecular weight bands highlighted with the arrow indicate BAI2. Note that molecular weights shift due to the loss of domains. The ΔTSP deletion mutant is the only construct that loses immunoreactivity to the anti-BAI2 antibody.

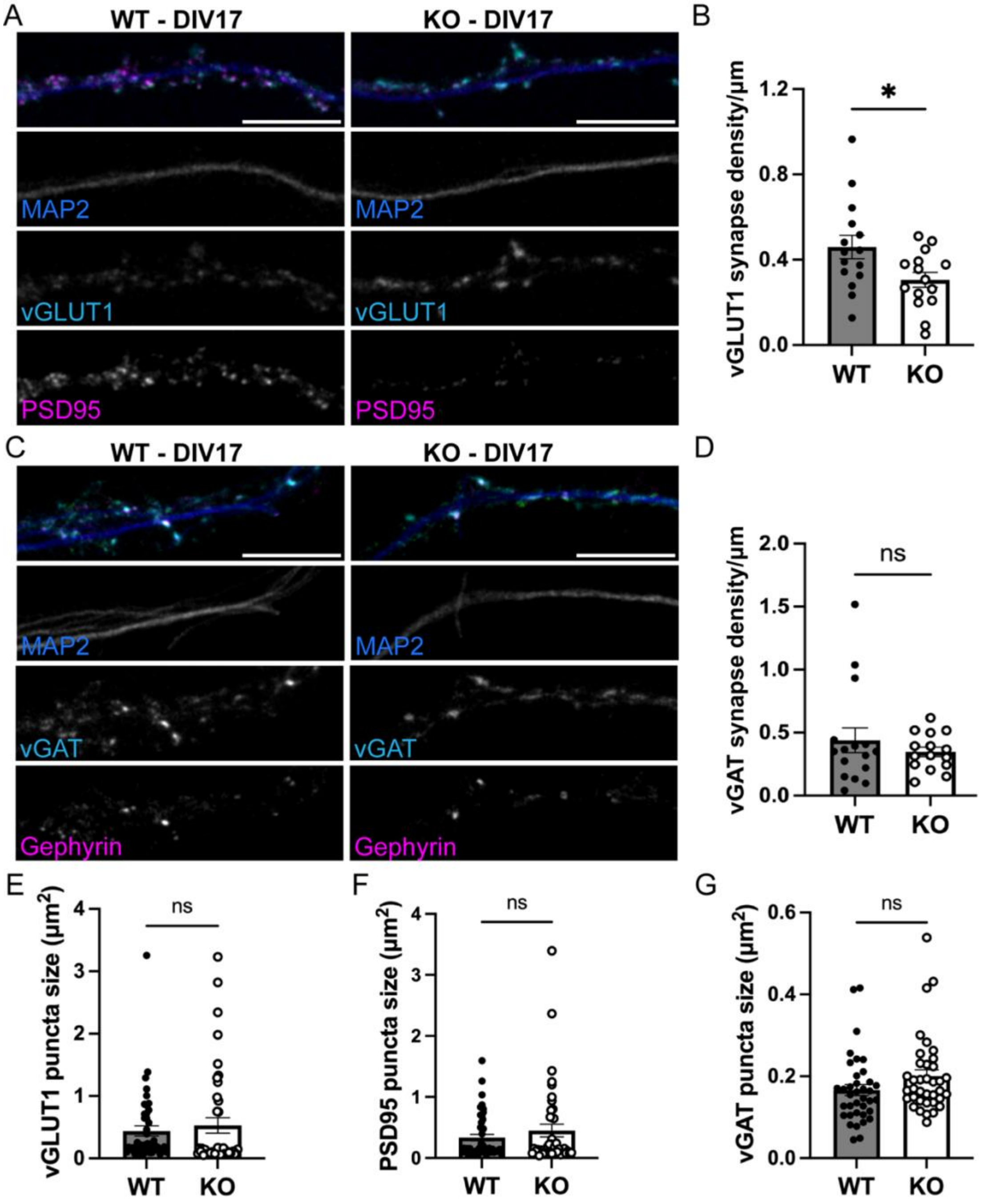


**Supplemental Figure 2. Loss of BAI2 results in reduced excitatory synapse density at DIV17**

(A, C) Representative images of hippocampal neurons from wild-type (WT) and *Bai2* knockout (KO) mice at DIV17 immunostained with presynaptic markers (vGLUT1 or vGAT) and postsynaptic markers (PSD95 or gephyrin). (B, D) Graphs depict density of vGLUT1-positive (B) and vGAT-positive synapses (D) at DIV17. (E, F, G) Graphs depict puncta size for PSD95 (E), vGLUT1 (F), and vGAT (G) at DIV17. Data represent mean±SEM. Statistics: unpaired t-test with or without Welch's correction, ns, not significant, $p^{*}<0.05$, $p^{**}<0.01$; n = 14-16 cells/group. Scale bar: 10 µm.
